# Chondro/osteoclasts and mast cells are co-villains in the joint destruction of rheumatoid arthritis

**DOI:** 10.1101/2023.01.04.522721

**Authors:** John McClure, Sheena F McClure

## Abstract

Chondro/osteoclasts and mast cells are cells of interest in the cartilage and bone destruction of joints affected by rheumatoid arthritis (RA). Both are major cellular components of the vascular synovial pannus proliferation characteristic of this disease. Chondroclasts degrade calcified cartilage and osteoclasts degrade bone tissue. Chondroclasts and osteoclasts are identical cell types and differentiate from monocyte precursors. Our studies show a close microanatomical relationship between these cells and new capillary formation (shown by the lectin *Psophocarpus tetragonolobus* – PTL-11) in the resorption sites of the mineralized tissues. Clast and mast cells express receptors for the lectin lPHA indicating beta1,6-acetylglucosaminal transferase V (GNTase V/MGAT5) activity providing a mechanism for neoangiogenesis. In addition to an angiogenetic function for mast cells it is probable that their products control monocyte differentiation and chondro/osteoclastogenesis.

## INTRODUCTION

In rheumatoid arthritis (RA) the synovial pannus is the hyperplastic and hypercellular synovial membrane which spreads over the articular surface of diarthrodial joints adhering to and, eventually, destroying the cartilage.

The constituent cells of the pannus have, over time, received varying degrees of attention being in and out of scientific fashion. In Gardner’s 1972 otherwise comprehensive book ‘The Pathology of Rheumatoid Arthritis’, two cells, the osteoclast and the mast cell, receive scant, if any, attention and yet are now considered to be major players. The osteoclast is not mentioned and the only reference to mast cells is the presence of granules in rheumatoid lung similar to the intracytoplasmic granules of mast cells (1).

Bilroth (more famous for surgical innovations) described multinucleated cells in synovial pannus in 1882. Perceptively he called these “bone breakers” (2). Thereafter osteoclasts went “off the radar” until rediscovery by Bromley and Woolley in 1984 (3). They described resorptive (clast) cells on calcified cartilage (chondroclasts) and subchondral bone (osteoclasts). Leison *et al* in 1988 (4) described the junction between cellular pannus and cartilage and bone as the erosive front with resorption bays typical of osteoclastic activity. Gravallese *et al* (5) in 1988 conclusively showed that chondroclasts and osteoclasts were identical with both showing the features characteristic of osteoclasts. A contemporary view is that the importance of osteoclasts in the pathogenesis of RA is such that they warrant being therapeutic targets (6).

Also in 1984 Bromley, Fisher and Woolley identified mast cells at sites of erosion of cartilage in RA. They were located at sites suggestive of newly formed pockets of calcified matrix erosion often in association with small blood vessels. Comparatively few mast cells were observed in pannus remote from the erosion front (7). Later Tetlow and Woolley (8) showed that mast cell activation in RA pannus was frequently associated with proinflammatory cytokine and metalloproteinase expression in neighbouring cells suggesting an important role for the mast cell in matrix degradation in microfoci of cartilage destruction. Most recently (2020) Min *et al* have pointed out that the mast cell proteases (eg tryptase) are well known angiogenetic factors in neoplasia. Mast cells also secrete various other proangiogenetic factors such as VEGF, basic fibroblast growth factors, TGF beta, TNF alpha and IL-8. They, therefore, suggested that the role of mast cells in angiogenesis in RA needs to be studied. Another recent concept is that mast cells in RA promote osteoclastogenesis by driving differentiation of monocyte precursors present in the synovial pannus (9).

In the present study the availability of complete joint samples (comprising articular surface, subchondral bone, pannus and portions of capsule) allowed us to conduct a detailed microanatomical study of mast cells, osteoclasts, blood vessels, their locations and relationships in rheumatoid arthritis.

## MATERIAL AND METHODS

As part of the joint replacement surgery for rheumatoid arthritis excision, samples became available for histopathological study. They consisted of joint surface with proximal bone tissue, synovial pannus and variable portions of the joint capsule. Specifically, there were three ulnar heads, two metacarpal heads and one radial head. The samples were derived from four male and two female patients with a mean age of 58 years (range 43 - 72).

All samples were fixed in formol-saline for a minimum of 48 hours and then decalcified in EDTA under x-ray control. Paraffin wax-embedded tissue blocks were prepared by conventional techniques. Thin sections were variously stained with haematoxylin and eosin (H & E), for mast cell tryptase and with the lectins lPHA and PTL-11 (10,11).

## RESULTS

Erosion of uncalcified cartilage, calcified cartilage and subjacent bone was by a vascular connective tissue (synovial pannus) of variable cellular composition and generally vascular. Pannus adherent to and degrading uncalcified cartilage contained blood vessels, macrophages and mast cells. Pannus degrading calcified cartilage typically had high concentrations of chondroclasts at the resorption zone. Closely associated with these were capillary blood vessels and mast cells many of which were degranulating. Resorption of the bone plate was by osteoclasts again with companion capillary vessel and mast cells in focal surface resorption pits. Resorption of osteonal and trabecular bone was by osteoclast tunnelling also with companion capillary vessels and mast cells. All these features were observable in H & E stained tissue sections.

Positive staining with the lectin lPHA was seen in type A synoviocytes of the pannus, in mast cells, in endothelial cells, in chondroclasts and osteoclasts. Positive staining with the lectin PTL-II was only ever observed in endothelial cells. Mast cells were universally demonstrated by positive staining for mast cell tryptase. In addition to the specific microanatomical locations described above, mast cells were randomly distributed in relatively small numbers in the more distal bone marrow.

## DISCUSSION

These observations confirm the close microanatomical association of mast cells, clast cells and blood vessels. This reinforces the concept that mast and clast cells are co-villains in the joint destruction of rheumatoid arthritis, as it particularly affects the layer of calcified cartilage and the adjacent subchondral bone. The degradation of uncalcified hyaline cartilage does not require this collaboration. In both locales the pannus shows a neovascularization indicated by positive staining by the lectin PTL-11. This confirms an O-glycosylation necessary for the formation of competent vascular tubules (12).

We applied the lectin lPHA to tissue sections in the present study since we have previously shown that clast and mast cells and endothelial cells stain positively with this lectin (10,11). The specific ligand is the β1-6 branching linkage in complex N-glycans and is formed by the action of N-acetylglucosaminyltransferase-V (GnTase V/MGAT5). This enzyme is overexpressed by carcinomas and chondrosarcoma and is believed to promote tumour growth and metastasis probably via angiogenesis (13). It has been postulated that MGAT5 selectively remodels the endothelial cell surface by the regulated binding of Galectin-1 (Gal1) which on recognition of complex N-glycans on VEGFR2 activates VEGF signalling. Also, in experimental animals with complete absence of MGAT5, N-glycans on VEGFR bind insufficient Gal-1 to promote angiogenesis and thus tumour growth decreases (10).

There is also substantial evidence linking mast cells and angiogenesis. Mast cells accumulate in a number of angiogenesis dependent processes such as wound healing and tumour growth. Several mast cell mediators (TNF, VEGF, FGF-2) are angiogenic. In addition, mast cell products such as tryptase can degrade connective tissue matrix providing space for neovascular sprouts (10).

That clast cells can drive angiogenesis is clearly important since a local, new and competent neovasculature is necessary for the effective removal of matrix breakdown products. Collaboration of clast and mast cells in neovascularization is clearly feasible. Expression of MGAT5 by endothelial cells may be an angiogenetic drive by an autocrine mechanism.

There is recent evidence of a more complex relationship between clast and mast cells. Histamine which is a major constituent of mast cells promotes osteoclastogenesis through the differential expression of histamine receptors on osteoclasts and osteoblasts (14,15) and stimulates osteoclast differentiation in monocytes (16). The osteoclastic mediators MdK and CXCL10 are released from mast cells in an oestrogen dependent manner via the oestrogen receptor alpha on mast cells (17). The relationship is a close one and is probably that of a master and servant. These findings provide a rationale for attempting anti-mast cell and anti-osteoclast therapies in the treatment of rheumatoid arthritis.

Tanaka (6) points out that radiographic studies have shown that bone erosion in RA begins at an early stage of disease and gradually (sometimes rapidly) exacerbates. The present study shows cortical and trabecular bone resorption without immediate contact with pannus and, therefore, at a distance to it making related bone structures vulnerable to breakdown.

**Figure 1.**
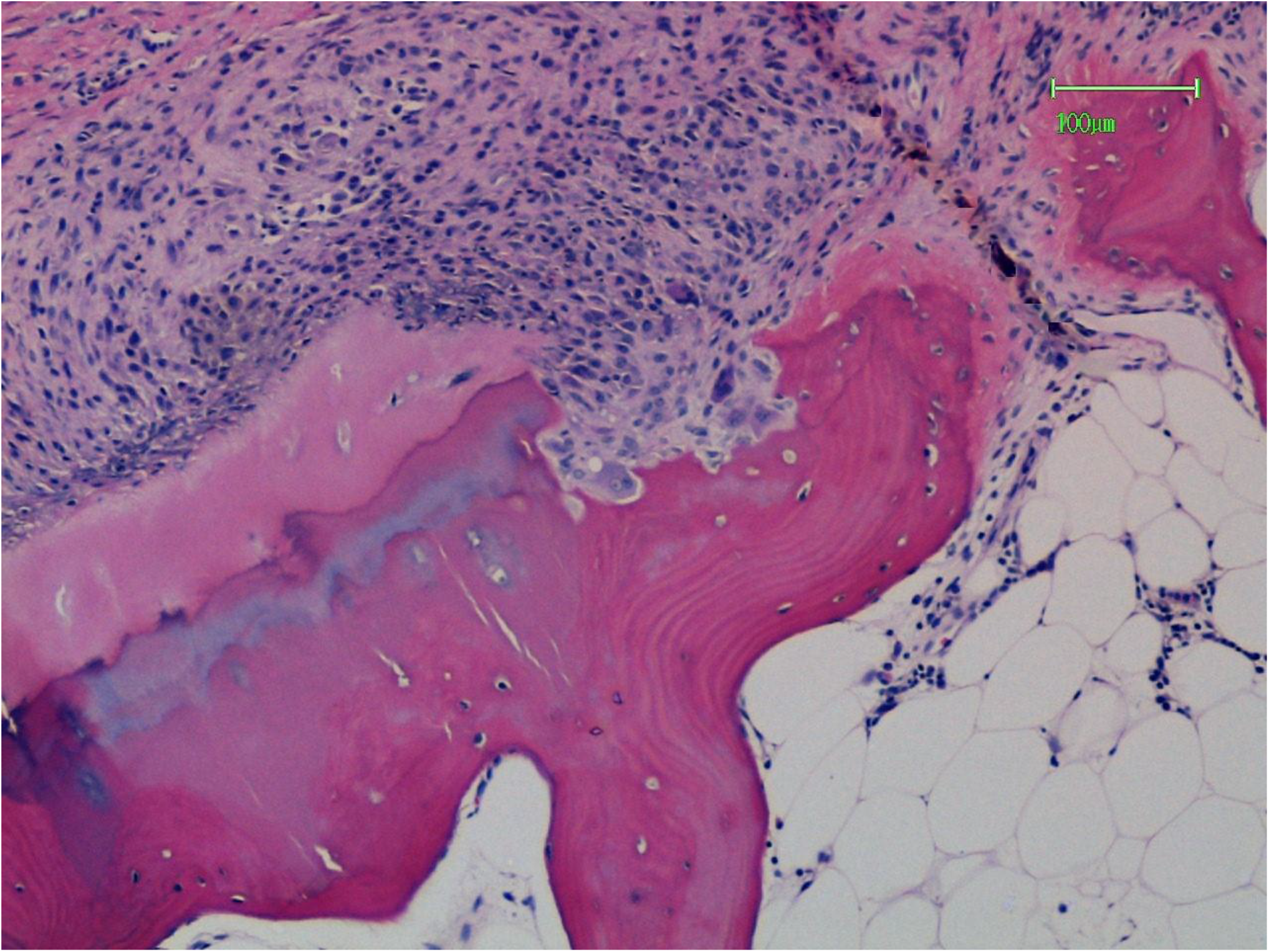
The richly cellular pannus has penetrated all layers of the articular cartilage into the subchondral bone plate. Erosion bays containing osteoclasts are present in the latter. Haematoxylin and Eosin.

**Figure 2.**
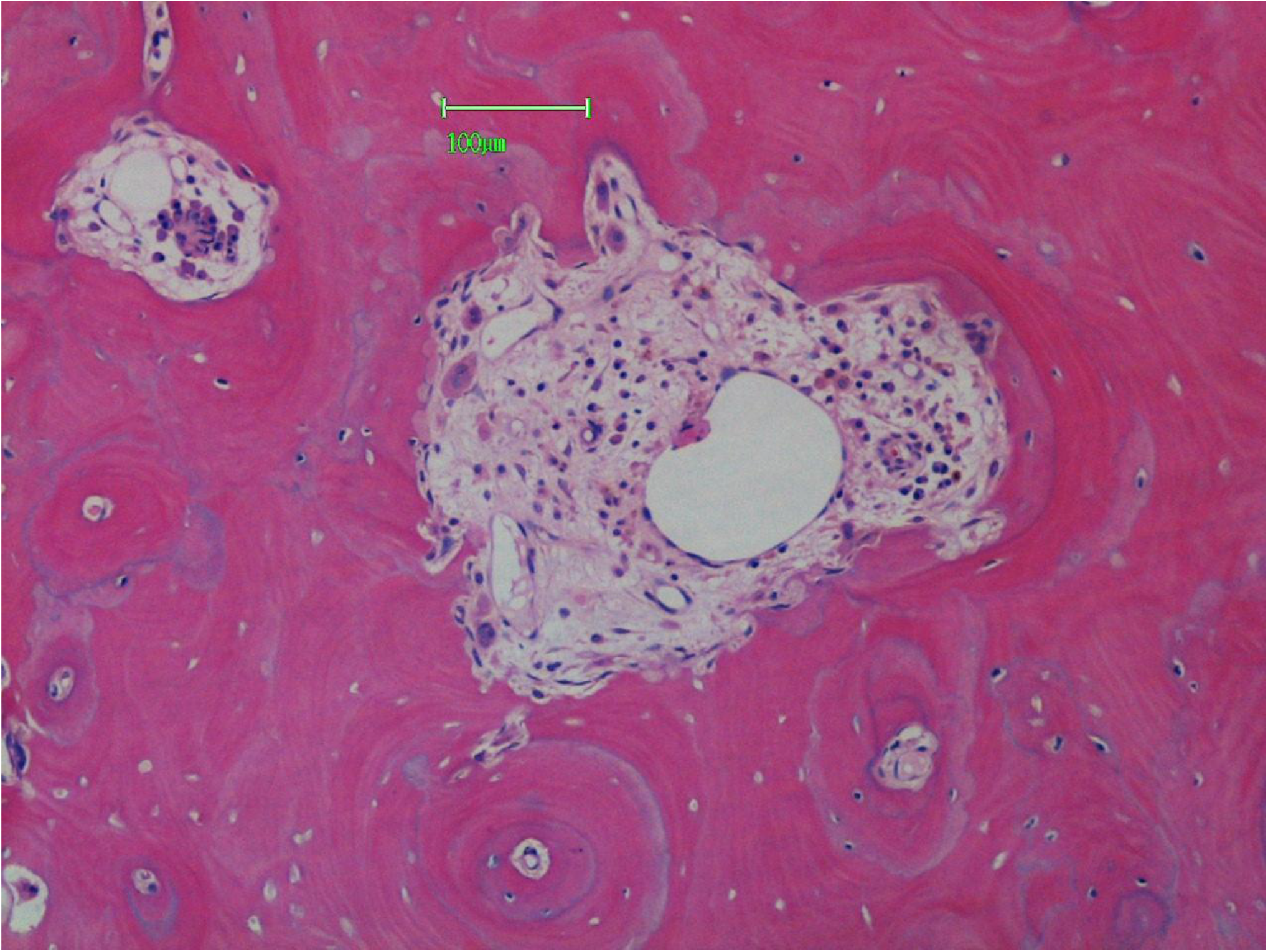
Erosion of osteonal bone by cutting cones (seen in cross-section) containing a nexus of osteoclasts, mast cells and capillaries. Haematoxylin and Eosin.

**Figure 3.**
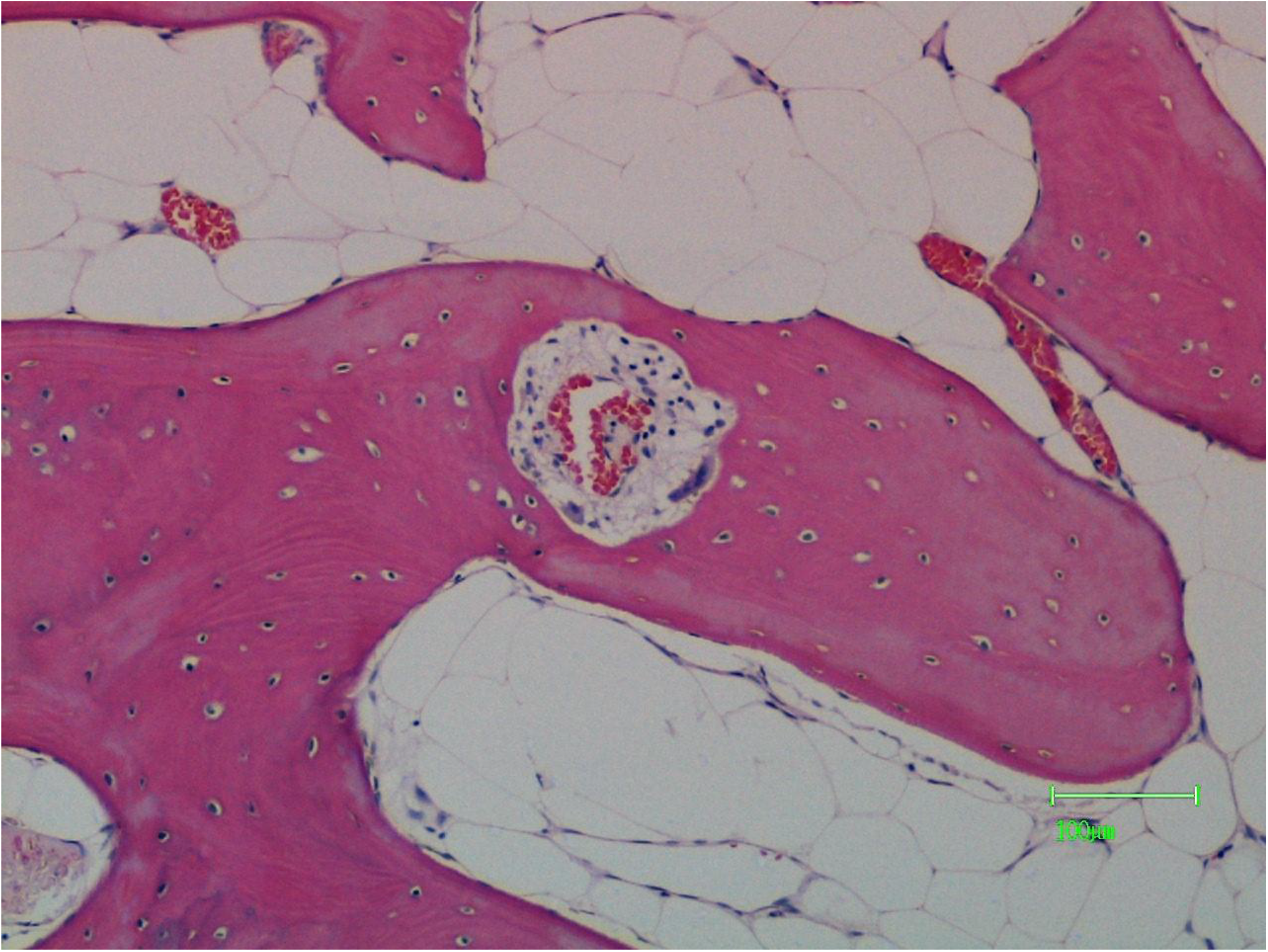
Tunnelling of trabecular bone by a cutting cone containing osteoclasts, mast cells and capillaries. Haematoxylin and Eosin.

**Figure 4.**
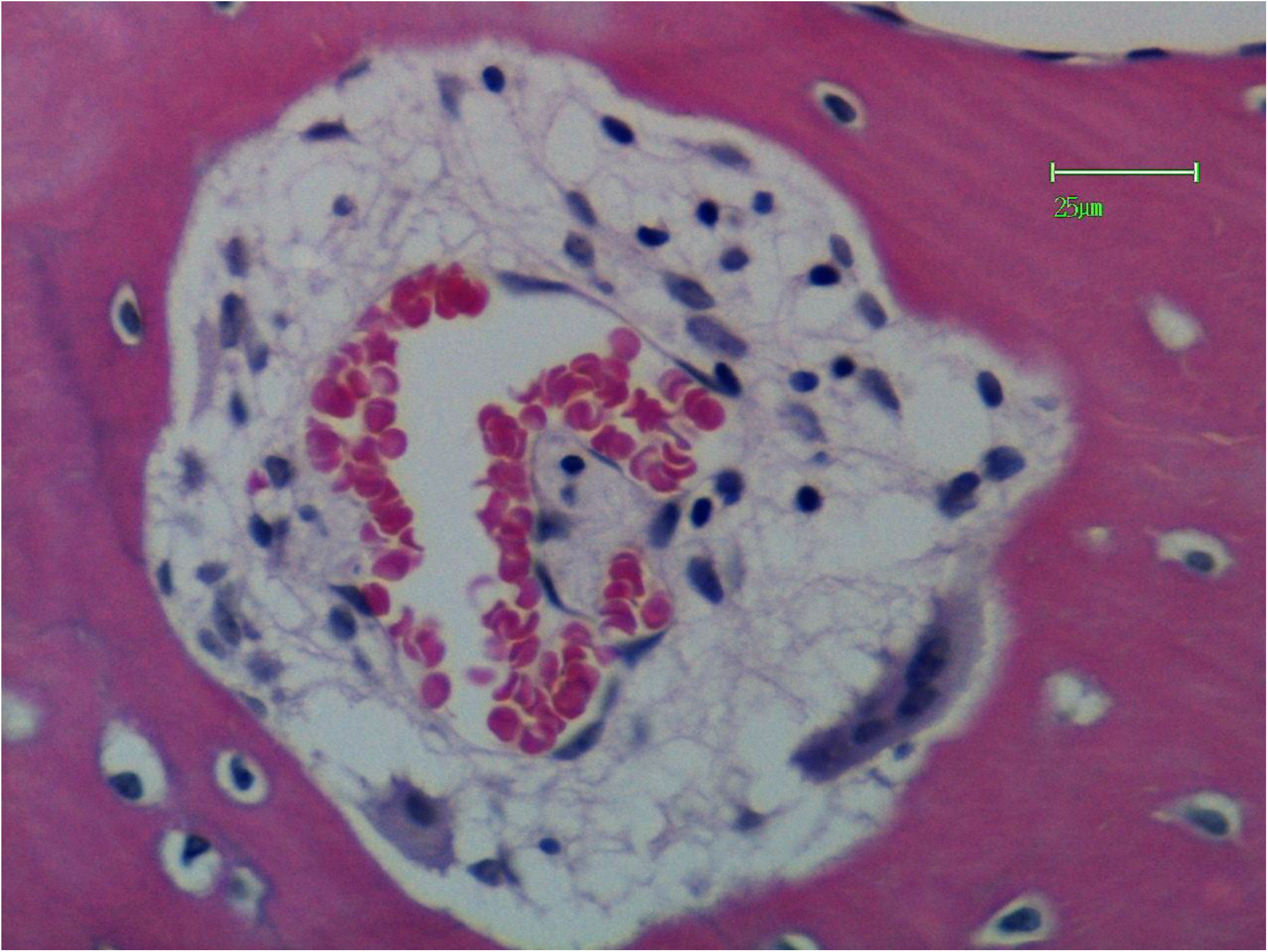
Higher magnification of the resorption tunnel shown in Figure 3. Haematoxylin and Eosin.

**Figure 5.**
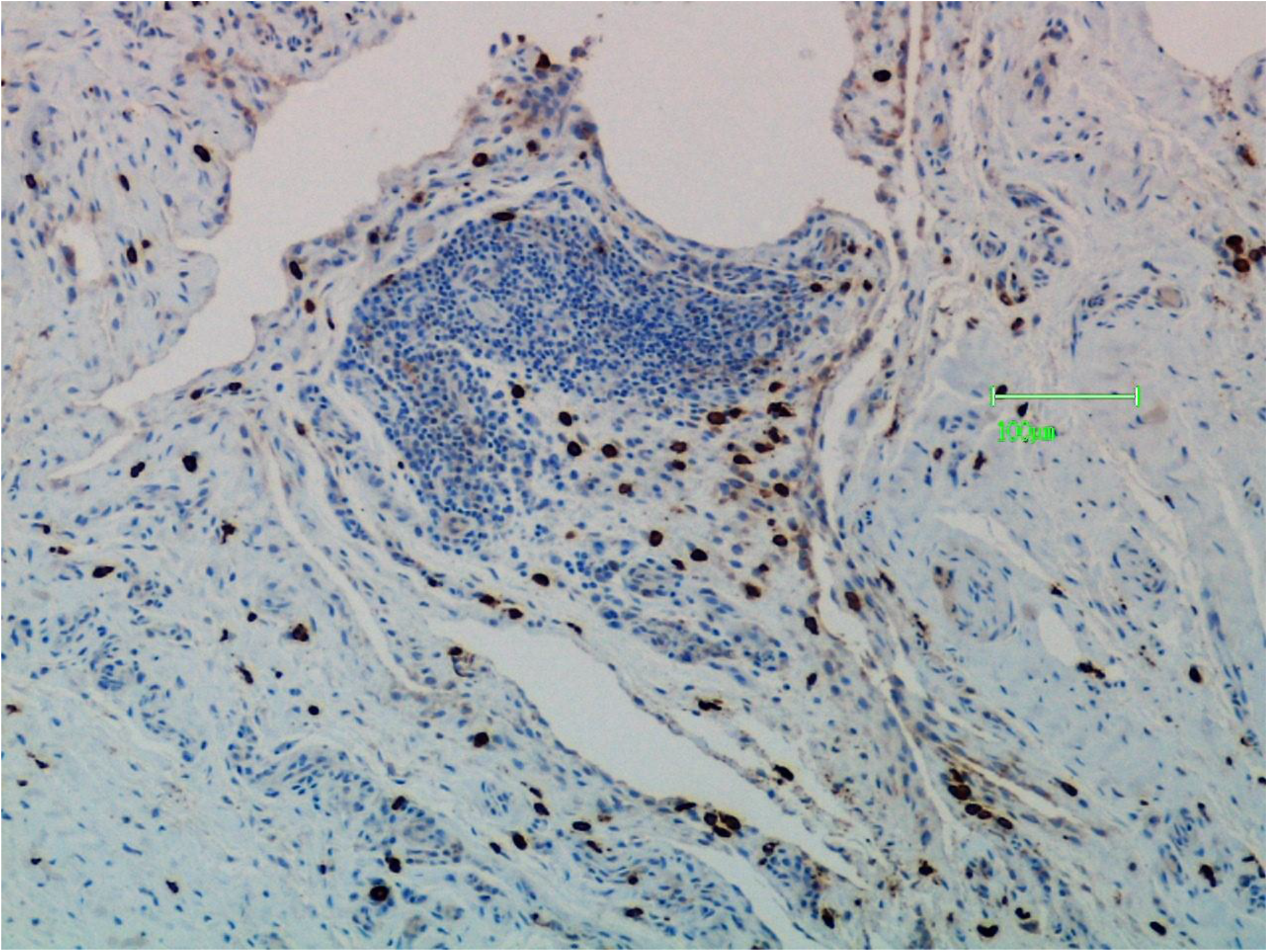
The mast cell population of the synovial pannus is demonstrated by staining for mast cell tryptase.

**Figure 6.**
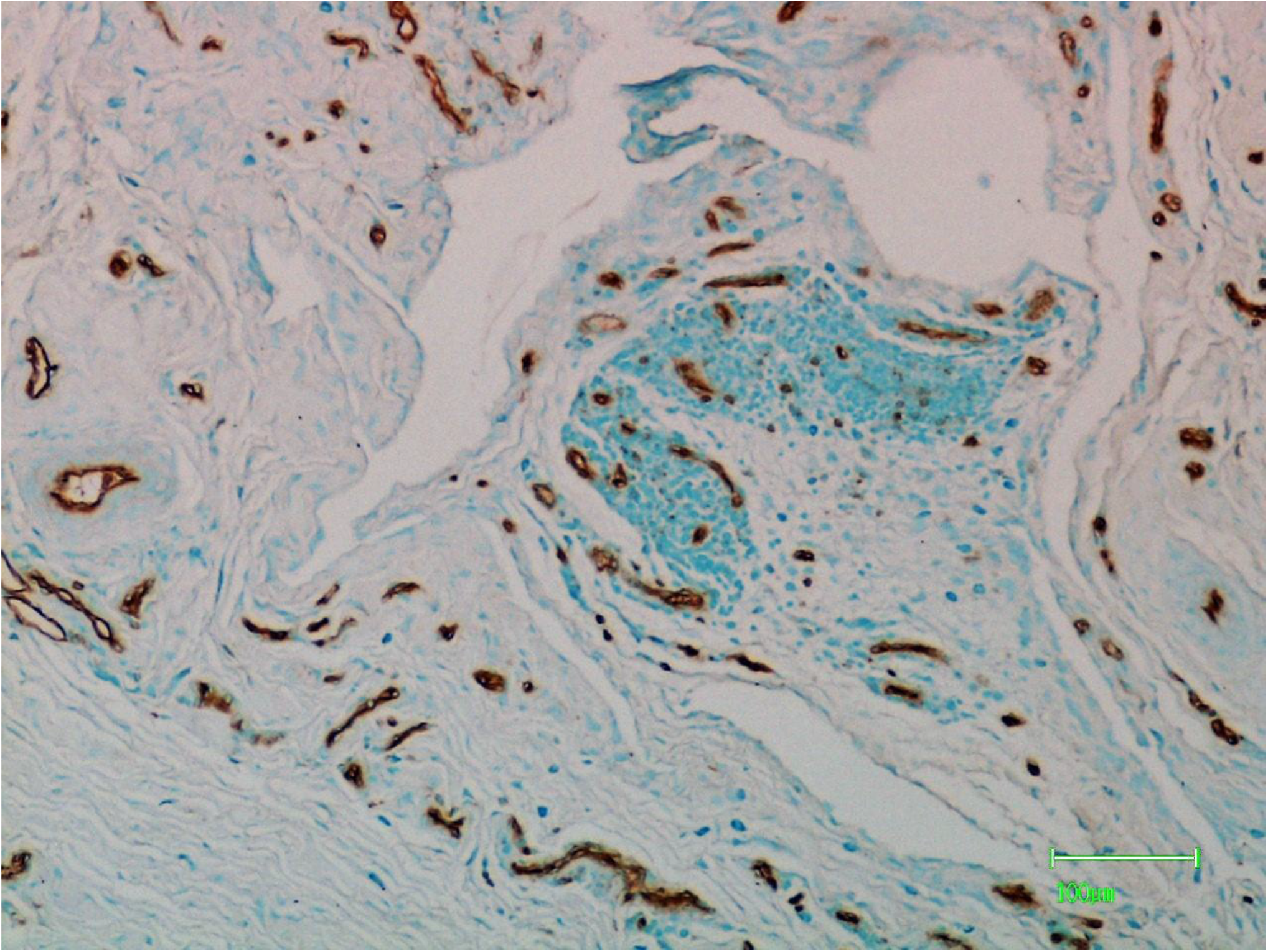
The parallel field of Figure 5 has been stained by the lectin PTL-11 to illustrate new blood vessels.

**Figure 7.**
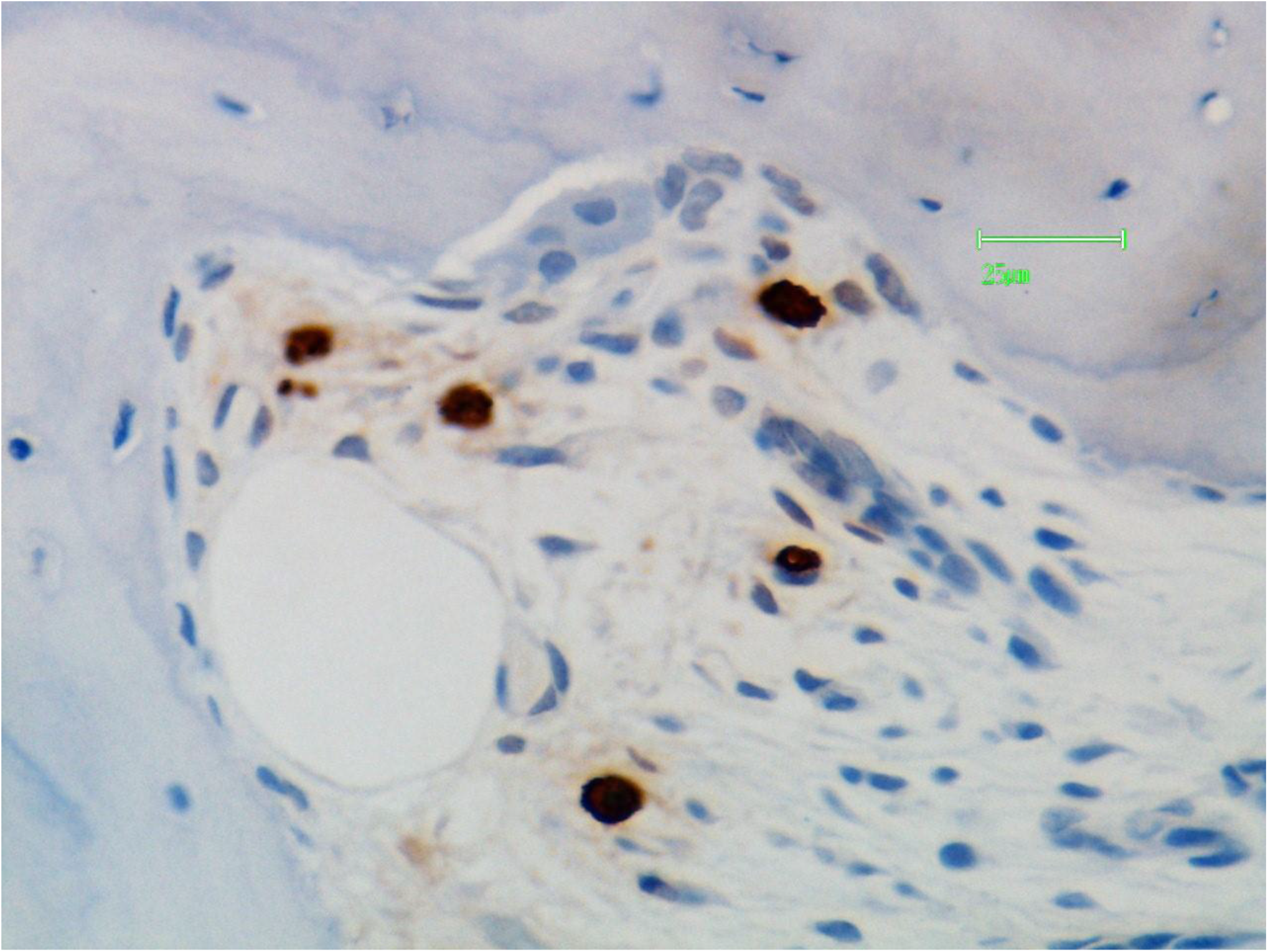
The contents of a resorption bay are osteoclasts, mast cells (stained for mast cell tryptase) and capillaries.

**Figure 8.**
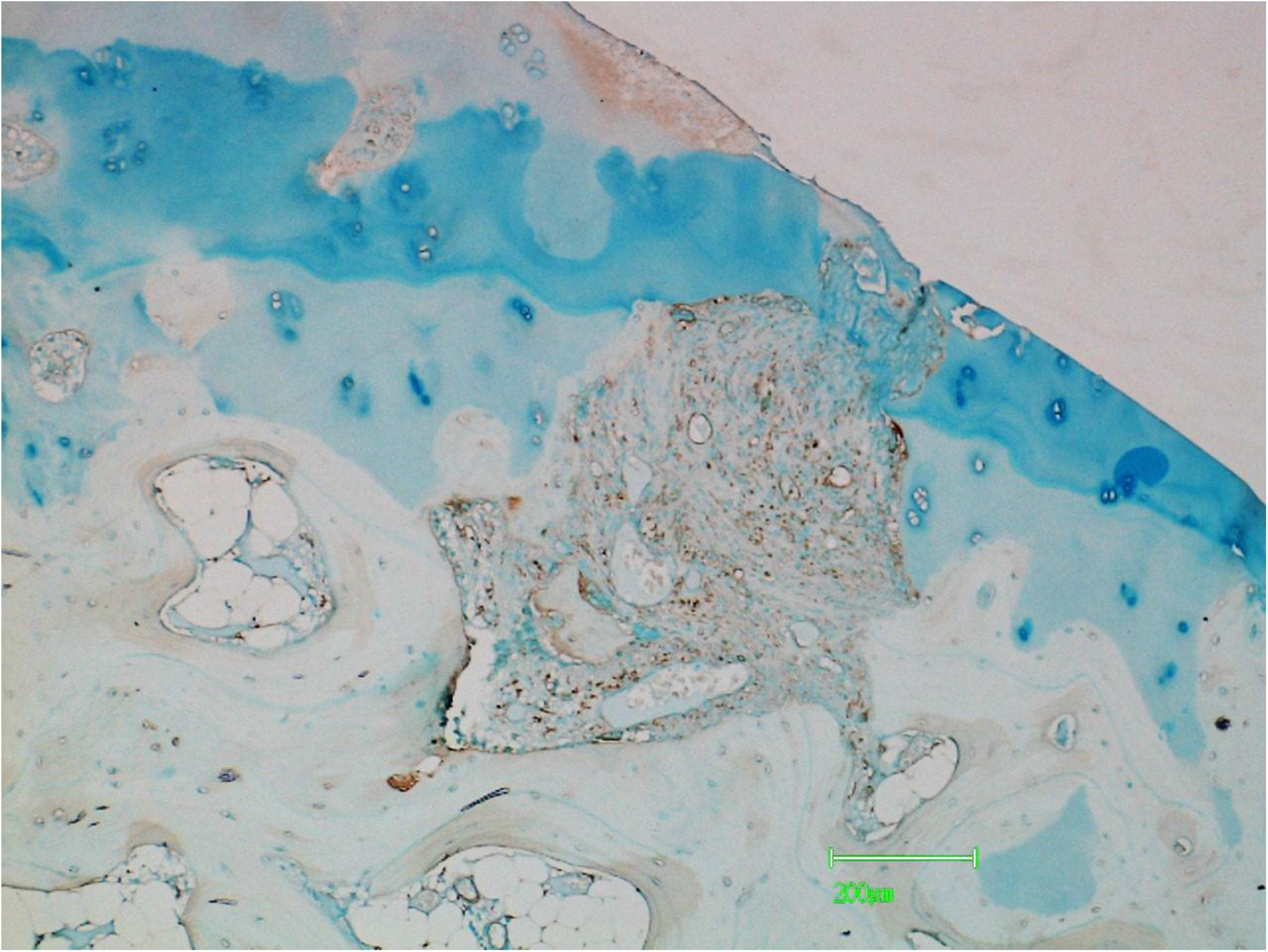
The synovial pannus has penetrated the uncalcified hyaline cartilage, tidemark, calcified cartilage and subchondral bone. Staining for the lectin lPHA produces positive reactions in chondroclasts, mast cells and blood vessels.

**Figure 9.**
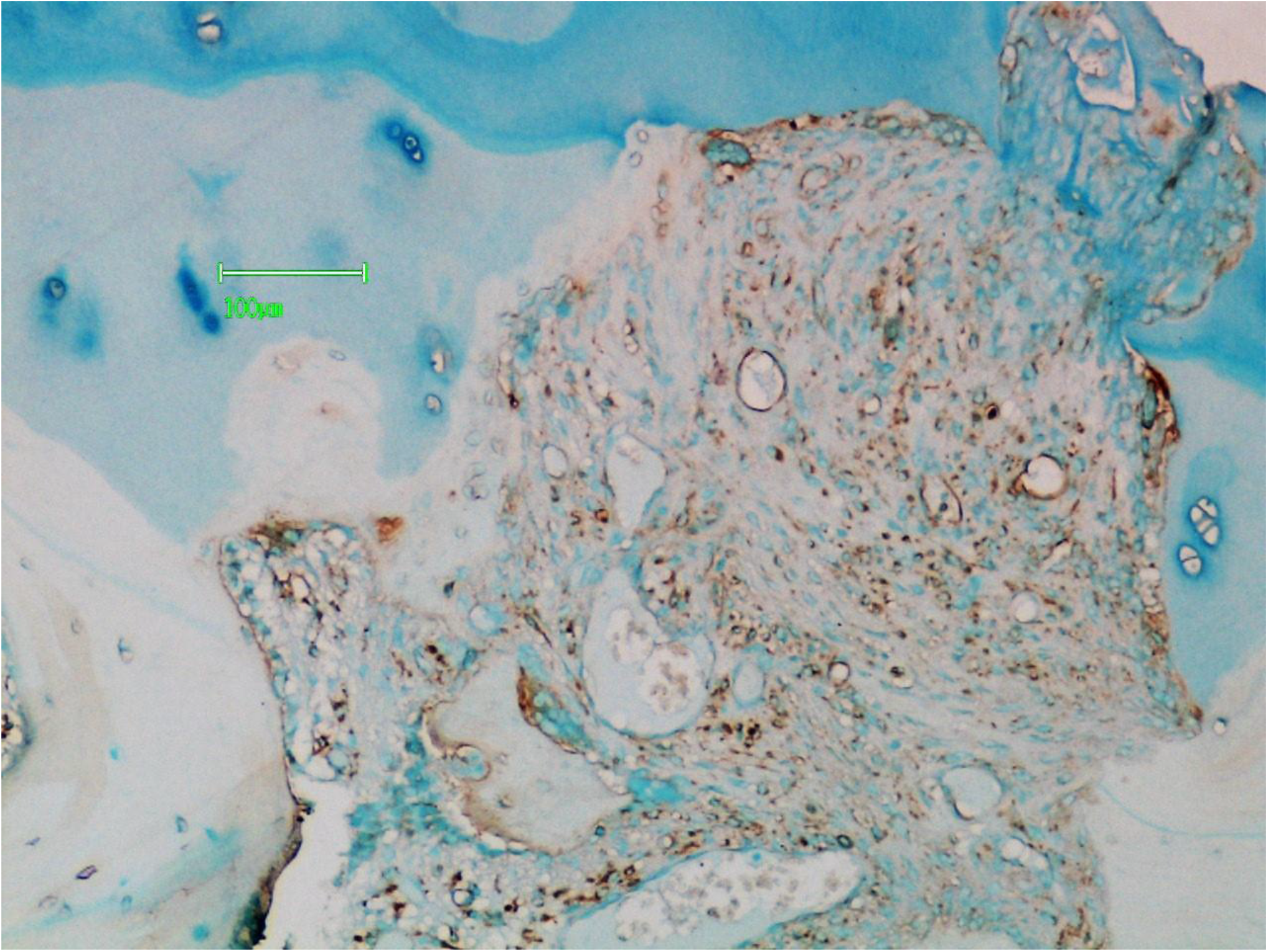
The same field as in Figure 8 at higher magnification showing the staining reactions in better definition.

**Figure 10.**
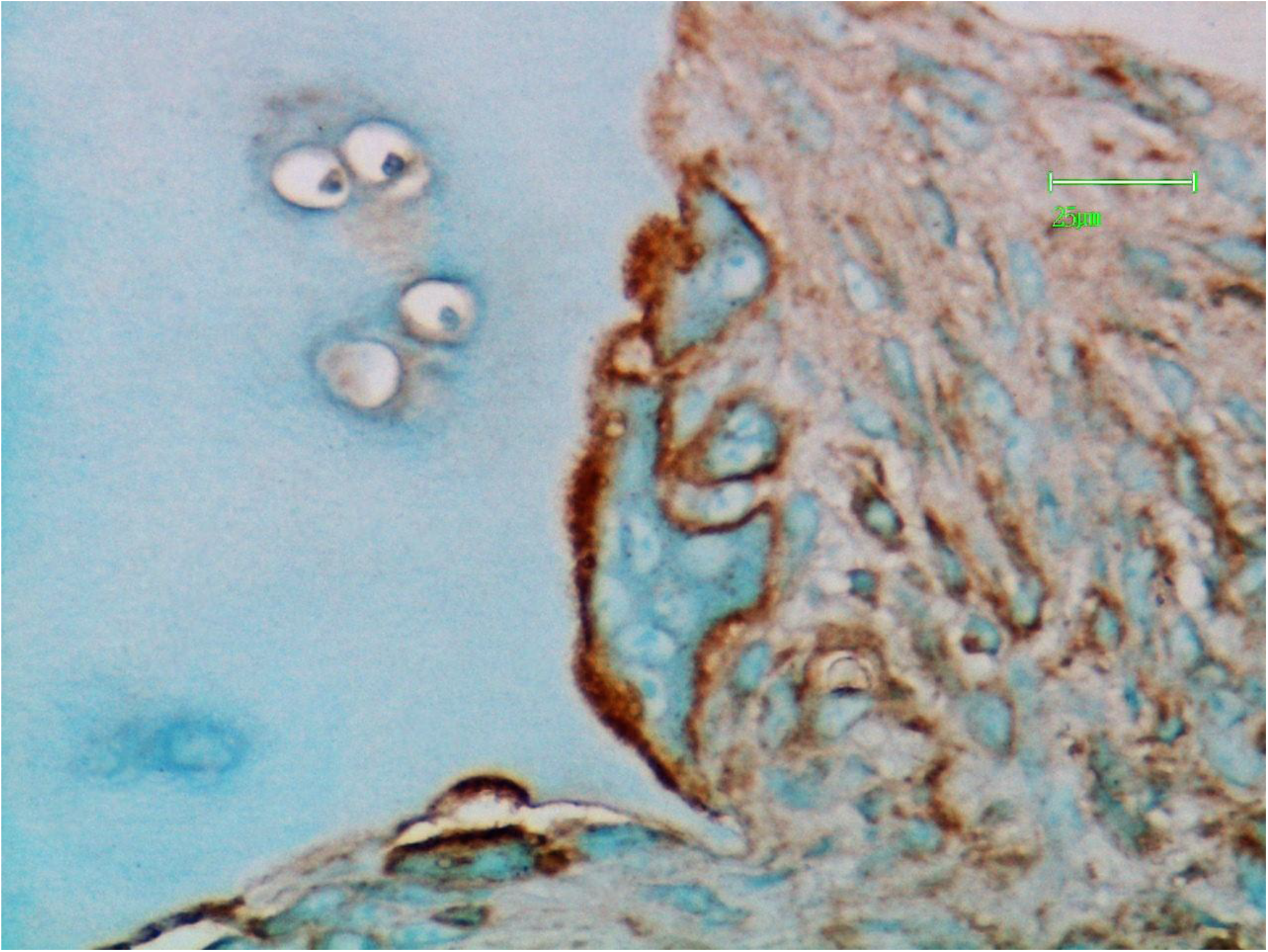
A chondroclast resorbing calcified cartilage has been stained for lPHA ligand. Positive reactions are membrane associated and particularly prominent on the ruffled border of the resorbing surface.

